# Analysis of Metabolic Network Disruption in Engineered Microbial Hosts due to Enzyme Promiscuity

**DOI:** 10.1101/2020.09.02.279539

**Authors:** Vladimir Porokhin, Sara A. Amin, Trevor B. Nicks, Venkatesh Endalur Gopinarayanan, Nikhil U. Nair, Soha Hassoun

## Abstract

**Background:** Increasing understanding of metabolic and regulatory networks underlying microbial physiology has enabled creation of progressively more complex synthetic biological systems for biochemical, biomedical, agricultural, and environmental applications. However, despite best efforts, confounding phenotypes still emerge from unforeseen interplay between biological parts, and the design of robust and modular biological systems remains elusive. Such interactions are difficult to predict when designing synthetic systems and may manifest during experimental testing as inefficiencies that need to be overcome. Despite advances in tools and methodologies for strain engineering, there remains a lack of tools that can systematically identify incompatibilities between the native metabolism of the host and its engineered modifications.

**Results:** Transforming organisms such as *Escherichia coli* into microbial factories is achieved via a number of engineering strategies, used individually or in combination, with the goal of maximizing the production of chosen target compounds. One technique relies on suppressing or overexpressing selected genes; another involves on introducing heterologous enzymes into a microbial host. These modifications steer mass flux towards the set of desired metabolites but may create unexpected interactions. In this work, we develop a computational method, termed <u>M</u>etabolic <u>D</u>isruption Work<u>flow</u> (*MDFlow*), for discovering interactions and network disruption arising from enzyme promiscuity – the ability of enzymes to act on a wide range of molecules that are structurally similar to their native substrates. We apply *MDFlow* to two experimentally verified cases where strains with essential genes knocked out are rescued by interactions resulting from overexpression of one or more other genes. We then apply *MDFlow* to predict and evaluate a number of putative promiscuous reactions that can interfere with two heterologous pathways designed for 3-hydroxypropic acid (3-HP) production.

**Conclusions:** Using *MDFlow*, we can identify putative enzyme promiscuity and the subsequent formation of unintended and undesirable byproducts that are not only disruptive to the host metabolism but also to the intended end-objective of high biosynthetic productivity and yield. In addition, we show how enzyme promiscuity can potentially be responsible for the adaptability of cells to the disruption of essential pathways in terms of biomass growth.

## 1. Introduction

Integrating heterologous synthesis pathways within microbial hosts has been instrumental in the biomanufacturing of industrial products such as biofuels, polymers, pharmaceuticals, therapeutics, flavors and chemical commodities [1–5]. One strategy to improve yield is to use well-established metabolic engineering techniques such as gene deletion, promoter engineering, media optimization, etc. [6]. Another strategy is to directly engineer enzymatic properties such as activity, selectivity, inhibition-resistance, and solubility [7]. Using one or more of these strategies has proven effective in development of strains with desired target yields, productivity, and titers.

Many such metabolic engineering strategies, however, yield unexpected enzyme-compound interactions. Some interactions can be beneficial for the survival of the host. For instance, Patrick et al. documented 41 rescue instances where the lethality of an essential protein deletion was suppressed by overexpression of one noncognate *E. coli* gene, attributing some of them to catalytic promiscuity and substrate ambiguity [8]. In other cases, beneficial interactions can come at a cost. The overexpression or knockout of enzymes can result in interactions that are disruptive for growth and maintenance by siphoning off key metabolic intermediates like pyruvate, acetyl-coA, and NADH. For example, while seeking to suppress lethality of inactivating the pyridoxal-5-phosphate (PLP) cofactor synthesis pathway, Kim et al. experimentally identified a four-step serendipitous pathway in *E. coli* that restored the strain’s ability to grow on glucose at the expense of consuming essential intermediates in the native serine biosynthetic pathway [9] and producing toxic byproducts [10].

The presence of high concentrations of heterologous enzymes and metabolites within microbial cells causes unexpected promiscuous interactions with host enzymes and metabolites. For example, the short-chain dehydrogenase YMR226C used to produce 3-HP in yeast is associated with 15 known substrates [11, 12]. Importantly, at high concentrations, promiscuous enzymatic reactions occur at noticeable levels. For example, in pathways intended for butanol production, the promiscuity of the bifunctional butyryl-coA dehydrogenase (*adhe2*) enzyme with substrate acetyl-coA often results in concomitant synthesis of ethanol with butanol [13–15]. In addition, despite efforts to reduce acetate production and force flux towards other metabolites by knocking out phosphate acetyltransferase (*pta*), acetate is still formed in these strains [14]. Similarly, lactate and acetate are produced at high levels, 0.27 g/L and 7.11 g/L respectively, even after lactate dehydrogenase (*ldh*) and *pta* were knocked out when producing 3-hydroxypropionic acid (3-HP) via the malonyl-CoA pathway in *E. coli* [16]. Liao and colleagues leveraged promiscuity of ketoacid decarboxylase (*KVD*) and alcohol dehydrogenase (*ADH*) enzymes to synthesize a spectrum of alcohols from branched-chain amino acid metabolic intermediates [17]. Yet, a caveat of this promiscuous activity is that no single alcohol can be made alone. That is, pyruvate itself is a ketoacid, which can be converted to ethanol by the same promiscuous activity of *KVD* and *ADH.* Thus, isobutanol synthesis is also coupled to ethanol synthesis due to enzyme promiscuity [18].

While ubiquitous [19–22] and often observed, effects of enzyme promiscuity on the host metabolic network are often ignored during design and only identified during experimental studies. Predicting such interactions early in the design cycle can yield improved design outcomes and reduce experimental efforts. The prediction of enzymatic products due to substrate promiscuity has mainly relied on hand-curated rules that capture well-known enzymatic transformations. For example, a list of 50 reaction rules, each associated with one or more reaction, was previously defined to explore novel synthesis pathways [23, 24]. Another set of rules was applied repetitively to generate novel synthesis [25] or degradation pathways [26]. Further use of such rules allowed the compilation of over 130,000 hypothetical enzymatic reactions that connect two or more KEGG metabolites [27], and the compilation of predicted metabolic products into databases such as MINEs [28]. MyCompoundID [29] utilizes a similar paradigm and generates products by the repeated application of addition or subtraction of common functional groups. BioTransformer [30] predicts derivatives by utilizing five separate prediction modules in concert with machine learning and a rule-based knowledge base. The PROXIMAL algorithm was utilized to create organism-specific <u>E</u>xtended <u>M</u>etabolic <u>M</u>odels (EMMs) that extend reference metabolic models catalogued in databases to include putative products due to promiscuous native enzymatic activities on native metabolites [31, 32]. PROXIMAL utilizes enzyme-specific reactant–product transformation patterns from the KEGG database [33] as a lookup table to predict products for query molecules [34]. Despite advances in predicting promiscuous products, however, these efforts have not been put forward in a systematic way to analyze metabolic network disruption in engineered microbial hosts due to enzyme promiscuity.

We develop in this paper a computational method, <u>M</u>etabolic <u>D</u>isruption Work<u>flow</u> (*MDFlow*), to analyze the disruptive impact of enzyme promiscuity in engineered microbial hosts in a systematic manner. We define *metabolic disruption* as changes in host metabolism due to enzyme-substrate interactions that may be due to gene overexpression or heterologous synthesis pathways, where such interactions do not exist neither in the wild-type chassis organism nor in the intended heterologous pathway. Accordingly, this definition encompasses all enzyme promiscuity that arises as a consequence of adding heterologous enzymes and their chemical products to a host microbe. Therefore, we have developed *MDFlow* to consider two different disruption scenarios. In “Scenario 1”, promiscuous activity is predicted in the context of overexpressed synthesis pathway enzymes, whether heterologous or native, acting promiscuously on native host metabolites. Meanwhile, in “Scenario 2,” predictions are made by assuming that native enzymes exhibit promiscuous interaction with synthesis pathway metabolites introduced with engineering changes. Of course, in a biological system, both scenarios would occur simultaneously to some degree, and even higher-order interactions would be possible (e.g., subsequent use of promiscuous reaction products as substrates for additional transformations).

To analyze such putative interactions, *MDFlow* relies on PROXIMAL, a tool for predicting byproducts resulting from promiscuous activities. Stoichiometrically balanced reactions are then derived based on substrate, product, and the reaction associated with the promiscuous enzymatic transformation pattern. To assess the disruption impact in a systematic fashion, the host metabolic network model is incrementally modified and evaluated after each change – first as is (to set a baseline), then augmented with the engineering strategy of interest, and lastly, with predicted byproducts and their associated reactions incorporated in the engineered network. Flux Balance Analysis (FBA) [35] is used to evaluate the impact of such changes on biomass growth rates or product yield. The results are then compared to expose the effects of both the deliberate and unexpected interactions.

We demonstrate the use of *MDFlow* to evaluate how engineered microbial hosts are impacted by enzyme promiscuity under different engineering strategies. For this work, we used *E. coli* as a microbial host, modeled using the *i*ML1515 and *i*ML1428 metabolic networks [36]. We demonstrate the formation of unexpected – and potentially disruptive – enzymatic transformations within the host under both scenarios.

We first focus on the rescue instances reported by Patrick et al. In that work, the authors set out to identify and categorize multifunctional genes that enabled cells’ adaptability to genetic lesions [8]. Within the Keio collection [37], the authors identified 107 single-gene knockout strains that were unable to grow on the M9-glucose medium. Patrick et al. found 21 strains that could be rescued via overexpression of one non-cognate gene, for a total of 41 unique suppression examples. Comparing structural superimpositions of deleted proteins and their suppressors, Patrick et al. observed significant structural homology between 6 enzyme pairs and attributed 11 examples to substrate ambiguity and catalytic promiscuity [8]. The fact that an essential gene deletion was suppressed by overexpression of another native gene and that enzyme/substrate non-specificity played a role in the rescue suggests that the responsible mechanism in many instances could have been a single enzymatic reaction mediated by promiscuous activity of overexpressed enzymes – that is, activity following Scenario 1. *MDFlow* is thus used with observations made by Patrick et al. to predict enzymatic reactions with a profound rescuing impact on host metabolism.

We then investigate how *MDFlow* can be applied to identify serendipitous multi-step interactions, similar to the pathway found by Kim et al. [9]. In that work, the authors identified a four-step chain of interactions that compensated for an essential gene knockout. The *pdxB* gene deletion disrupted the PLP synthesis pathway, resulting in its inability to grow on M9-glucose at 37 °C. Simultaneous overexpression of seven other genes, however, rescued the strain. Using genetic complementation experiments, the authors were able to separate those genes into groups and describe one of the rescuing pathways in detail. The serendipitous pathway was found to bypass the knocked-out enzyme in the PLP pathway by diverting flux from serine biosynthesis. The interactions comprising this pathway were catalyzed by three enzymes – two of which exhibited either promiscuity (ThrB) or broad specificity (LtaE). The function of the third enzyme (YeaB) was unknown, and one of the interactions appeared to be non-enzymatic. Because *MDFlow* anticipates interactions between native metabolites and engineered modifications in the scope of the entire cell, with one or more genes being overexpressed simultaneously, it can be used to identify similar multi-step interactions where a native pathway contributes metabolites to an unexpected serendipitous process. Similarly, to the rescue instances considered by Patrick et al., such interactions are simulated by *MDFlow* as a part of Scenario 1.

Lastly, we evaluate the potential disruption of two metabolic pathways that when added to *E. coli* yield 3-hydroxypropionic acid (3-HP), an important precursor metabolite to useful derivatives such as acrylic acid, 1,3-propanediol, and malonic acid [38–40]. In this instance, *MDFlow* is used to analyze both disruption scenarios. We demonstrate that the result of both scenarios is the production of unexpected – and potentially disruptive – interactions within the host.

This work is novel as it is the first to systematically investigate effects of heterologous and native enzyme promiscuity on host metabolism and its consequences on biocatalysts. Our method serves as the first computational tool that can assist metabolic engineers in (1) identifying sources of unexpected byproducts, (2) assessing the consequences of metabolic engineering on the host, and (3) quantifying pathway-host incompatibility using metabolic network disruption. Outcomes from this work will aid in future studies to design robust systems with more predictable behaviors and improved desired product yield.

## 2. Methods

*MDFlow* (**Figure 1**) relies on FBA [35] to evaluate the yield or growth rate of the host and on PROXIMAL [34] for promiscuous products prediction. The workflow begins by considering the “initial” model that reflects the selected host and its environmental conditions. An “engineered” model is then created by introducing engineered modifications of interest. Lastly, the workflow constructs a set of interactions that can arise due to enzyme promiscuity and incorporates them into the engineered model, creating a new “disrupted” model. At every step, the behavior of the model is evaluated using FBA to determine the effect of each incremental change under different engineered conditions.

**Figure 1.**
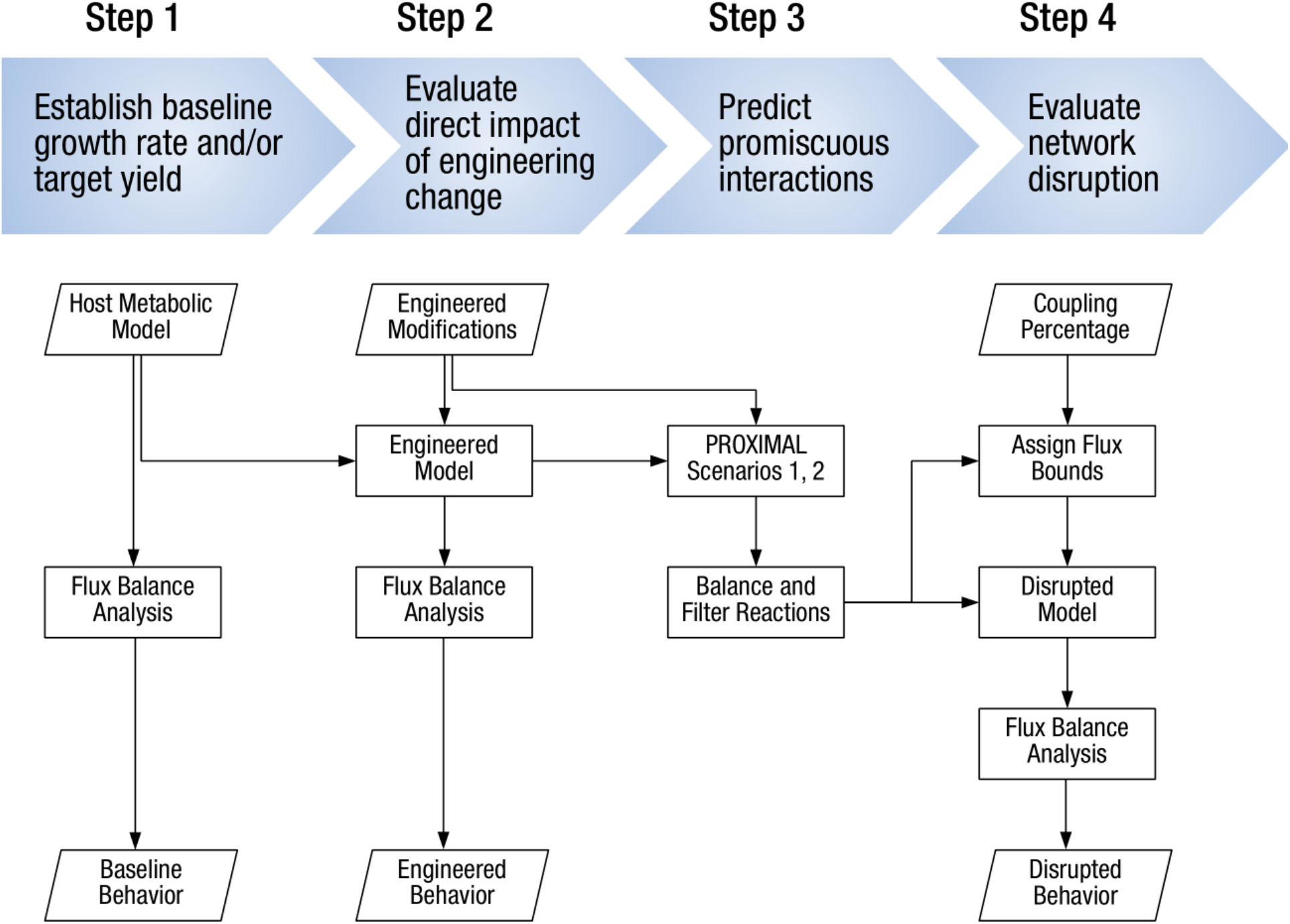
An overview of the four-step process used by *MDFlow* to identify and evaluate byproducts formed due to enzymes promiscuity for Scenarios 1 and 2. The original host metabolic model is progressively augmented with engineered modifications and predicted interactions. The updated models are evaluated using FBA at different stages.

### 2.1. Step 1 – Establish the baseline growth rate and/or target yield using FBA

To model the *E. coli* metabolic network, we build upon the *i*ML1515 model published by Monk et al. [36]. When evaluating gene knockout modifications, we use a derivative of that model, *i*ML1428, that offers improved accuracy for such experiments. We use the conditions suggested by the authors of *i*ML1515 for constraint-based modeling – the lower bounds of all exchange reactions were set to zero, except for glucose, oxygen, and all inorganic ions: the lower bound for each of those reactions was set to −10, −20, and −1000 mmol gDW^−1^ h^−1^, respectively. With the constraints configured for aerobic growth on glucose, we evaluate the baseline growth rate, and target metabolite yield (if applicable), of the host using FBA. Then, we set the lower bound on biomass growth to be equal to 10 % of that amount to reflect the survival needs of the host.

### 2.2. Step 2 – Evaluate direct impact of the engineering change using FBA

We implement intended changes for metabolic engineering in the context of addition or removal of reactions and metabolites in the network. To construct a synthesis pathway for transforming a metabolite within a host into a target compound, we add each synthesis step to the stochiometric matrix (S-matrix) of the metabolic model as a new reaction, along with a demand reaction for the target metabolite. To model a gene knockout, we enact the effect of deleting the gene from the model by setting flux to zero for all inactive reactions, identified based on their individual gene-protein-reaction (GPR) rules [41]. The new “engineered” model is then evaluated using FBA to demonstrate the impact of introduced changes. At this stage, it is important to verify that the model’s prediction reflects expectations or experimental data (e.g., lethal knockdowns or an expected yield).

### 2.3. Step 3 – Predict promiscuous interactions using PROXIMAL

The modified model from the previous step is used to predict promiscuous byproducts for Scenarios 1 and 2. Each added reaction is assumed reversible unless indicated otherwise. For each scenario, PROXIMAL first creates a lookup table of all known biotransformation operators based on catalogued reactions within the model. Given a query molecule, PROXIMAL first identifies operators in the lookup tables that can act on the molecule. Then, the query molecule is mapped to possible byproducts. An operator acts on a query molecule if its reaction center and its nearest neighbor atom(s) exactly match those of the native substrate, as encoded in the lookup tables. Each potential byproduct is reported in the form of a mol file, which is then used to identify if the potential byproduct is a known metabolite. The mol file is matched to either a metabolite in the model, a KEGG ID, or a PubChem ID using InChIKeys [42] generated by an open-source chemical toolbox RDKit [43].

For Scenario 1, biotransformation operators are derived from the overexpressed enzymes along the synthesis pathway and applied to all native metabolites in the model. For Scenario 2, biotransformations are derived from native host enzymes within the model and applied to pathway metabolites in the engineered pathway. Applying PROXIMAL operators results in a list of byproducts for each scenario. For each predicted byproduct, we develop a new balanced enzymatic reaction based on the catalyzing enzyme’s reaction pattern. A reaction template with suitable cofactors is obtained from reaction(s) associated with each enzymatic biotransformation. A reaction is balanced when the number of atoms for each element on the reactant side matches those on the product side. Reactions are verified to be balanced using ChemPy [44]. If a reaction is not balanced, it is discarded and not considered for further analysis as it violates the assumptions of FBA.

### 2.4. Step 4 - Evaluate network disruption using FBA

To predict the effect of promiscuity, the engineered model network is augmented with balanced promiscuous reactions, thereby creating a “disrupted” model. The metabolites in new reactions are either mapped to existing metabolites in the model or added to the model along with the corresponding exchange reactions allowing unlimited export (but not import) of a metabolite from the host. Each predicted reaction is first added to the engineered model separately. Using FBA, the flux range of each promiscuous reaction is calculated by minimizing and maximizing the flux through it in both forward and reverse directions (*i.e. v*_fwd_min_, *v*_fwd_max_, *v*_rev_min_, *v*_rev_max_). The added reactions are either new to the model or form biotransformation routes that are catalyzed differently than those already in the model.

To model promiscuous activity, we assume a non-zero flux through the added reaction (*v*_fwd_added_, *v*_rev_added_) based on a pre-defined percentage *p*, referred to as a *coupling percentage,* of the calculated maximum flux. As in case of all reactions when performing FBA, the absolute maximum for any reaction is set at ±1,000 mmol/h/gDW. The constraints imposed on the reaction can be described by the following inequalities – the first one pertains to the forward flux through the reaction, while the second describes the reverse flux:

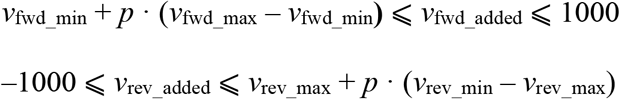

Once constraints are set, the metabolic network disruption caused by the developed reactions can be evaluated. The disrupted yield value is calculated using FBA and compared with the value of the yield of the undisrupted engineered model. The direction of each reaction is then fixed to the one that caused maximum disruption. For the purposes of subsequent analysis, reactions are considered irreversible and their individual flux is constrained by only one of the two inequalities. Disruption in yield is then evaluated under various coupling percentages, with a random subset of developed reactions added to the undisrupted model. The results are then placed on a scatter plot relating the extent of metabolic disruption to the intensity of promiscuous activity for visualization purposes.

## 3. Results

### 3.1. *MDFlow* explains how suppressors can rescue growth of essential gene deletions

We applied *MDFlow* to the 41 unique knockout-suppressor pairs, reported by Patrick et al. where a single essential gene deletion was suppressed by overexpression of one noncognate gene in *E. coli.* To establish the baseline for validating rescue, we first determined if the deletion of each of the 20 essential genes responsible for the 41 pairs arrests the growth of the strain. To this end, we performed gene knockouts in *i*ML1428, a model derived from the *i*ML1515 model that offers enhanced accuracy for gene knockout predictions, and we measured the biomass growth rate reported by FBA. Out of the 20 lethal knockouts reported by the authors, we were able to confirm lack of growth for 9 of them, accounting for 15 out of the 41 unique knockout-suppressor pairs. Since we cannot computationally validate rescue without first establishing the lethality of the genetic lesion, we only focused on those 15 cases where the deletion led to no biomass. For each of the suppressor genes, we used PROXIMAL to generate promiscuous reactions due to Scenario 1 type interactions (e.g., overexpressed gene acting on native metabolites) and then applied FBA to determine their individual effect on the biomass, resulting in the complete recovery of 6 strains (*ΔilvE, ΔglnA, ΔilvD, ΔhisH, ΔpabA,* and *ΔilvA*). In the majority of the recovered cases, a promiscuous reaction replicated the deleted essential reaction either exactly (e.g., *ΔilvA/tdcB, ΔpabA/pabB,* and *ΔhisH/hisF*) or with different cofactors (e.g., *ΔilvE/avtA* and *ΔglnA/asnB).* Both *pabA/pabB* and *hisF/hisF* form well-characterized heterodimeric enzymes. The larger subunits (PabB and HisF) of both enzymes have shown activity in the absence of the smaller subunits (PabA and HisH), which supports our predictions that PabB and HisF could execute compensating reactions in the *ΔpabA and ΔhisH* strains, respectively [45, 46]. Meanwhile, in the *ΔilvA/tdcB* case, the two proteins are known to be isozymes of one another, thus having the same function [8]. Therefore, there is biological justification for the rescuer enzyme’s potential ability to exactly replicate the reactions lost by the deletion. From the perspective of our method, both the deleted gene and its suppressor had the same Enzyme Commission (EC) numbers, which made them both interchangeable for the purposes of transformation pattern extraction. In the other 3 case sets, however, the outcome appears to be due to enzyme promiscuity – although the first two or three EC groups matched in two instances, the deleted protein and its replacement were not identical enzymes in all three. For *ΔglnA/asnB*, the promiscuous activity is responsible for creating a reaction edge between L-glutamate and L-glutamine, which appears to be the mechanism of recovery. For *ΔilvE/avtA,* the promiscuous reaction allows the production of L-isoleucine, which is essential for survival – as double mutants (*ΔilvE ΔavtA*) are known to require supplementation of both, isoleucine and valine, for growth [47]. Notably, the deletion of *ilvE* also caused the loss of an L-valine-producing reaction in the model, however, its essentiality was not confirmed through FBA. For *ΔilvD/avtA*, no single predicted reaction was responsible for growth recovery, however, including various combinations of two predicted reactions demonstrated rescuing effect. A summary of multicopy suppression results can be found in **Table 1** along with example compensating reactions predicted for each case. A complete set of predicted reactions is provided in **Supplementary File 1**.

**Table 1:** Confirmed-lethal single-gene deletion strains from Patrick et al. [8] with the corresponding multicopy suppressors (MS) and representative subsets of predicted compensating reactions. Biomass growth rate in the disrupted network computed using FBA is presented as a percentage of the wild type strain growth rate.

| Deletion | Deficient Reactions | MS | MS Biomass | Compensating Reactions |
| --- | --- | --- | --- | --- |
| *ΔilvA*<br>(4.3.1.19) | L-threonine → 2-oxobutanoate + $\text{NH}_4^+$ | *tdcB*<br>(4.3.1.17, 4.3.1.19) | 100.00% | L-threonine → 2-oxobutanoate + $\text{NH}_4^+$ |
| | | | 100.47% | L-*allo*-threonine → 2-oxobutanoate + $\text{NH}_4^+$ |
| *ΔilvE*<br>(2.6.1.42) | (S)-3-methyl-2-oxopentanoate + L-glutamate → 2-oxoglutarate + L-isoleucine<br>3-methyl-2-oxobutanoate + L-glutamate → 2-oxoglutarate + L-valine | *avtA*<br>(2.6.1.66) | 100.00% | (S)-3-methyl-2-oxopentanoate + L-alanine → L-isoleucine + pyruvate |
| | | | 100.79% | 3 3-methyl-2-oxobutanoate + 2 L-alanine → 2 L-isoleucine + 3 pyruvate |
| *ΔglnA*<br>(6.3.1.2) | ATP + L-glutamate + $\text{NH}_4^+$ → ADP + L-glutamine + $\text{H}^+$ + $\text{PO}_4^{3-}$ | *asnB*<br>(6.3.5.4) | 100.94% | AMP + L-asparagine + L-glutamate + $\text{P}_2\text{O}_7^{4-}$ → L-aspartate + ATP + L-glutamine + $\text{H}_2\text{O}$ |
| | | | 101.26% | L-glutamate + $\text{NH}_4^+$ → L-glutamine + $\text{H}_2\text{O}$ |
| | | | 99.38% | AMP + L-asparagine + D-glucose 1-phosphate + L-glutamate → ADP-glucose + L-aspartate + L-glutamine + $\text{H}_2\text{O}$ |
|  |  |  | (and others) |  |
| *ΔilvD*<br>(4.2.1.9) | (R)-2,3-dihydroxy-3-methylbutanoate → 3-methyl-2-oxobutanoate + $\text{H}_2\text{O}$<br>(R)-2,3-dihydroxy-3-methylpentanoate → (S)-3-methyl-2-oxopentanoate + $\text{H}_2\text{O}$ | *avtA*<br>(2.6.1.66) | 100.23% | 2 Pyruvate + 3 L-valine → 2 (S)-3-methyl-2-oxopentanoate + 3 L-alanine<br>2 4-methyl-2-oxopentanoate + 3 L-alanine → 2 pyruvate + 3 L-valine |
| | | | 100.23% | 3 3-methyl-2-oxobutanoate + 2 L-alanine → 2 L-isoleucine + 3 pyruvate<br>2 L-leucine + 3 Pyruvate → 3 3-methyl-2-oxobutanoate + 2 L-alanine |
|  |  |  | (other combinations may be possible) |  |
| *ΔhisH*<br>(4.3.2.10, 3.5.1.2) | L-glutamine + phosphoribulosylformimino-AICAR-phosphate → AICAR + erythro-imidazole-glycerol-phosphate + L-glutamate + $\text{H}^+$ | *hisF*<br>(4.3.2.10) | 100.02% | L-glutamine + phosphoribulosylformimino-AICAR-phosphate → AICAR + erythro-imidazole-glycerol-phosphate + L-glutamate |
| | | | 100.02% | L-glutamine + phosphoribulosylformiminoaicar-phosphate → AICAR + erythro-imidazole-glycerol-phosphate + L-glutamate |
| | | | 100.05% | L-glutamine + 2 phosphoribulosylformimino-AICAR-phosphate → (S)-2-hydroxyglutarate + 2 AICAR + 2 erythro-imidazole-glycerol-phosphate |
|  |  |  | (and others) |  |
| *ΔpabA* | | *pabB* | 100.00% | chorismate + L-glutamine → 4-amino-4-deoxychorismate + L-glutamate |
| (2.6.1.85) | chorismate + L-glutamine → 4-amino-4-deoxychorismate + L-glutamate | (2.6.1.85) | 100.00% | 2 chorismate + L-glutamine → 2 4-amino-4-deoxychorismate + (S)-2-hydroxyglutarate |
|  |  |  | 100.00% | chorismate + L-glutamine → 4-amino-4-deoxychorismate + O-acetyl-L-serine |
|  |  |  | 100.00% | L-asparagine + chorismate → 4-amino-4-deoxychorismate + L-aspartate |
|  |  |  | 100.00% | L-glutamine + isochorismate → 4-amino-4-deoxychorismate + L-glutamate |
|  |  |  | 100.00% | L-glutamine + prephenate → 4-amino-4-deoxychorismate + L-glutamate |

### 3.2. *MDFlow* can identify multi-step bypasses to essential gene functions

We considered the role of promiscuous interactions in the serendipitous pathway shown in Figure 1 that recovers the host’s ability to grow on glucose after its PLP pathway is disrupted due to the deletion of *pdxB.* The new pathway (**Figure 2**) bypasses the lesion by diverting a metabolite (3-phosphohydroxypyruvate, 3-PHP) from the serine production pathway and converting it to a metabolite (L-4-phosphohydroxythreonine, 4-PHT) downstream the PLP synthesis pathway via a series of 4 *ad-hoc* reactions [9]. Assuming the reactions are due to promiscuity, discovering the first and/or the last step in this pathway entails predicting interactions between the candidate enzymes (i.e., native enzymes, with the exception of PdxB, with amplified promiscuity due to overexpression) and all metabolites present in the model. Then, after extending the model with the potential promiscuous products, interactions can be predicted again to discover the next steps. Repeating this process up to four times, in principle, would allow the extraction of the entire pathway. Unfortunately, this approach would be computationally intensive because when the pathway is unknown, it amounts to an exhaustive breadth-first enumeration of all possible promiscuous reaction sequences. However, given the serendipitous PLP pathway, we can prune search space to exactly one candidate (since each reaction is already known) to quickly determine if the exhaustive search would eventually find it. Using *MDFlow* configured in this way, we were able to rediscover 3 out of the 4 reactions along this novel pathway. The elusive step, the LtaE-catalyzed transformation of glycolaldehyde to L-4-hydroxythreonine (4-HT), was not predicted due to the lack of an appropriate reaction pattern in PROXIMAL’s operator lookup table. This observation highlights an important shortcoming of rule-based methods such as BioTransformer [30] and PROXIMAL [34]: their metabolism predictions are limited to the finite set of biotransformation rules encoded in their knowledgebase. Nonetheless, the results for this pathway and multicopy suppressors from [8] show that our technique can be used to discover experimentally validated unexpected interactions that lead to the survival of the host.

**Figure 2:**
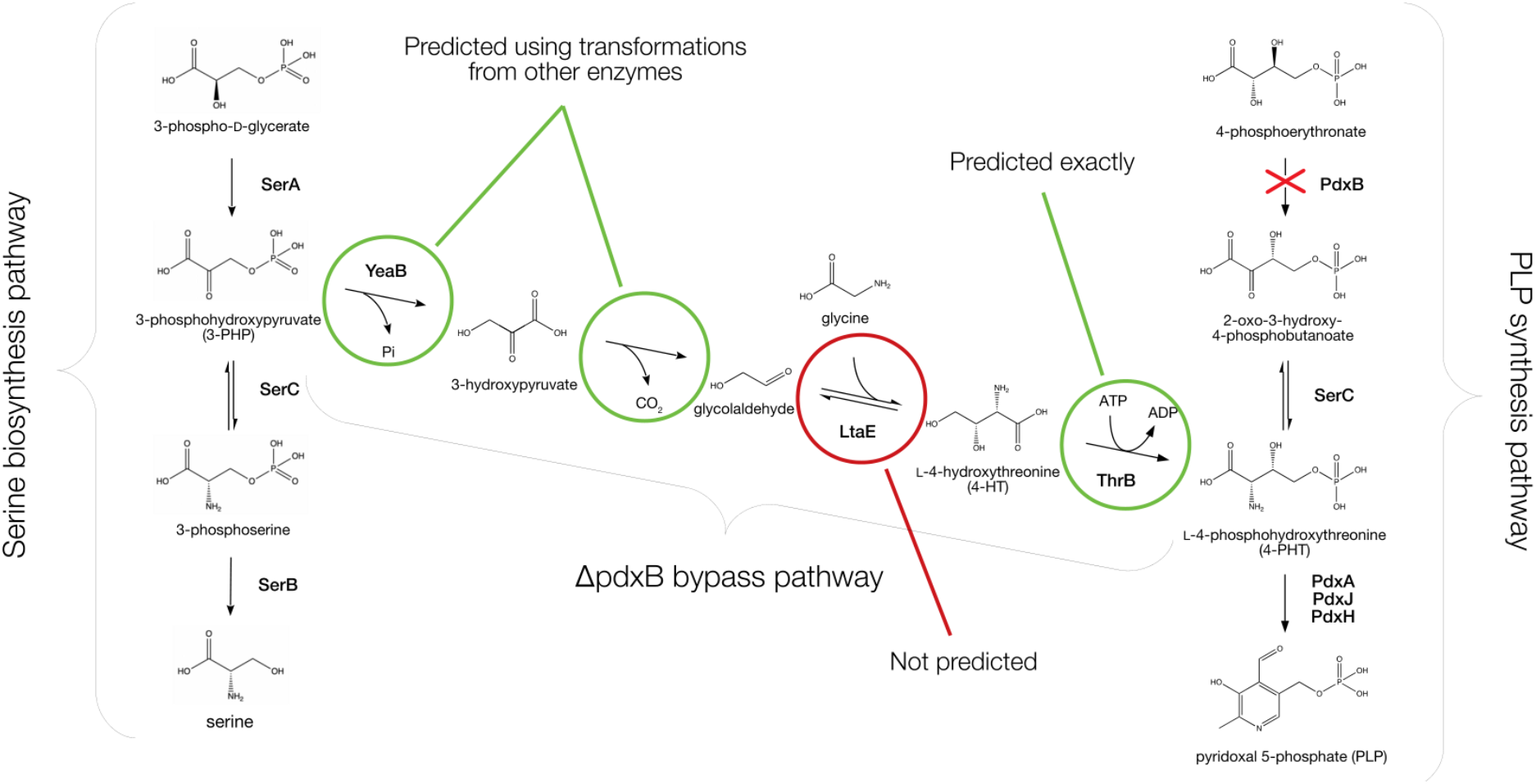
Serendipitous pathway discovered by Kim et al. that bypasses the deletion of an intermediate gene *pdxB* in the native PLP pathway by siphoning off material from the serine biosynthesis pathway. Circles highlight reaction steps that were predicted – or not predicted – by PROXIMAL. Pathway layout for the drawing was adapted from the authors’ original paper.

To further illustrate the utility of *MDFlow* in predicting multi-step pathways, we used *MDFlow* to discover two-step promiscuous pathways in iML1515 that may compensate for the deletion of *pdxB.* Such a two-step pathway transforms a metabolite within iML1515 to one of the following metabolites: 2-oxo-3-hydroxy-4-phosphobutanoate, 4-hydroxy-L-threonine, pyridoxal phosphate. These three metabolites are downstream from 4-Phosphoerythronate. We retained pathways that had intermediates with known PubChem IDs. We further filtered these results based on thermodynamic feasibility of the pathways as estimated by eQuilibrator [48–51]. There were 21 such feasible two-step pathways. There were 13 and 8 pathways that terminated on pyridoxal phosphate and 2-oxo-3-hydroxy-4-phosphobutanoate, respectively. There were none that terminated on 4-hydroxy-L-threonine. As the *i*ML1428 model does not capture the lethality of knocking out *pdxB,* we were unable to evaluate the impact of these two-step pathways on growth rates using FBA. A complete set of predicted two-step pathways is provided in **Supplementary File 2**.

### 3.3. *MDFlow* predicts metabolic disruption during 3-hydroxypropionic acid (3-HP) biosynthesis adversely affects yield

We used our workflow with two different synthetic 3-HP pathways as an example of application to metabolic engineering. Several groups have reported production of 3-HP in various organisms [52, 53], including *E. coli* [54, 55]. Using *E. coli* as a host, we first added a two-step 3-HP synthesis pathway (**Figure 3A** and **3B**) catalyzed by malonyl-CoA reductase (MCR) [56] and 3-hydroxy acid dehydrogenase (YdfG). Then, separately, we added a three-step 3-HP pathway comprising of *panD, gabT,* and *YdfG.* Assuming no disruption, the maximum theoretical yield of 3-HP was estimated to be 1.62 mol/mol of glucose for the first pathway and 1.66 mol/mol for the second. After the addition of a synthesis pathway to *E. coli,* our workflow utilized PROXIMAL to predict derivatives for both Scenarios 1 and 2. For both Scenarios, identities of predicted derivatives were looked up in either *i*ML1515, KEGG, or PubChem. As not all predicted derivatives were identifiable through this method, we considered only derivatives that are documented in at least one of the three databases. Example reactions for each Scenarios 1 and 2 are shown in **Figure 3, panels C** and **D**, respectively. Other predicted reactions are given in **Supplementary File 3**.

**Figure 3:**
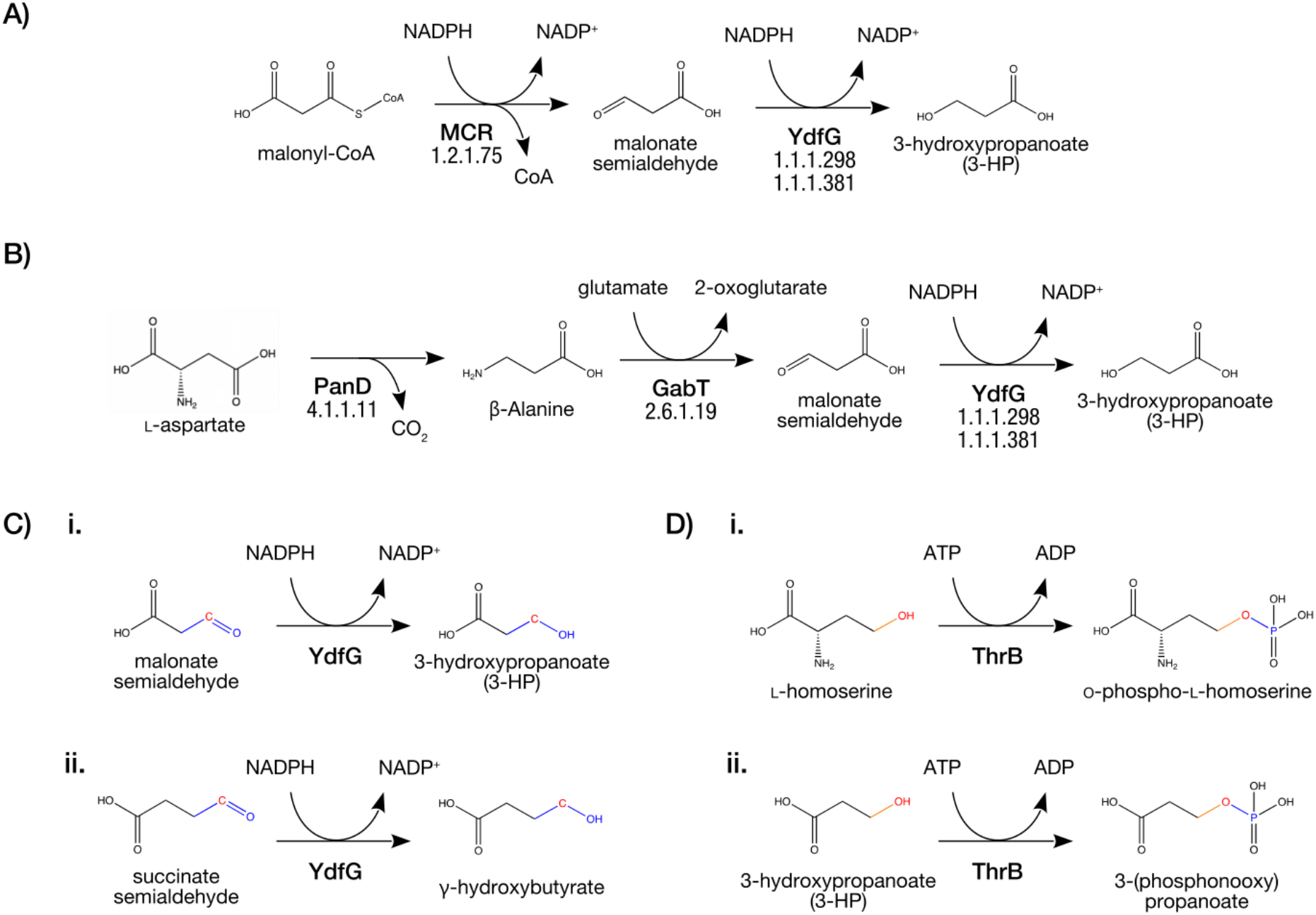
Overview of the two heterologous 3-HP pathways integrated into the *E. coli* model and the method used to construct putative promiscuous reactions for each scenario. (A) Pathway 1, which converts malonyl-CoA into 3-HP via two reactions catalyzed by malonyl-CoA reductase (MCR) and YdfG. (B) Pathway 2 that produces 3-HP from L-aspartate using PanD and GabT in addition to YdfG. Developed reactions examples of Scenario 1 (C) and Scenario 2 (D). Both panels (C) and (D) are divided in three sections: i. the native reaction catalyzed by the potentially promiscuous enzyme, ii. the RDM pattern showing the rction center in red where the biotransformation occurs, and iii. the developed balanced reaction indicating the reactants, products, and the promiscuous enzyme.

Balanced predicted reactions due to each scenario were first evaluated using FBA to determine the range of flux densities they can potentially sustain in the context of the *E. coli* metabolic model. The model was then augmented with all predicted reactions, along with the 3-HP synthesis pathway. We then assumed that each of the predicted reactions exhibits a certain minimum activity, measured as a fraction of its maximum possible flux. We refer to this fraction as the *coupling percentage* of a given reaction.

As not all enzymes and metabolites in *E. coli* are present in high concentrations, not all predicted promiscuous reactions occur at appreciable levels to disrupt the metabolic network. To address this issue, we performed randomized probabilistic analyses. For each scenario, we assumed that only a portion of the predicted promiscuous reactions act on their target molecules. First, a random mean coupling percentage *p*_mean_ was sampled from a normal distribution with *μ* = 1 % and *σ* = *μ* / 3. The standard deviation was chosen in such way to place 99.7 % of the samples within a range strictly above 0 %, and in the remaining 0.3 % of the cases, the coupling percentage was resampled repeatedly to obtain a value above 0 %, still. Then, we randomly selected 10 % of the developed reactions to exhibit promiscuous activity. For each of the selected reactions, a coupling percentage was chosen randomly from another normal distribution with *μ* = *p*mean and *σ* = *μ* / 3. The selected coupling percentage was then used to set the minimum required flux for the reaction according to one of the inequalities presented later in the methods section, while the lower and upper bounds of all other (non-selected) developed reactions were set to zero. 10,000 FBA runs were performed, each time selecting a different set of promiscuous reactions and re-sampling the mean and individual reaction coupling percentages. In each run, the FBA objective was to maximize the yield of 3-HP while maintaining biomass growth of at least 10 % of wildtype. The analysis was also repeated assuming 25 % and 50 % of the developed reactions to have promiscuous activity. Additionally, another run was performed, where the total number of active reactions was fixed at 10 and each Scenario was either allocated the entire set (10) or half of it (5). This was done to account for that fact that Scenario 2 reactions were much more plentiful than Scenario 1 reactions. Avoiding overrepresentation of the former reactions enables qualitative comparison of the two scenarios.

The results in **Figure 4 (A, B, C, and D)** for both pathways and scenarios exhibit similar trends. Higher disruption tends to be correlated with more active reactions and with higher coupling percentages. As the mean coupling percentage increases, so does the disruption, and the relationship is almost linear in all but Pathway 1, Scenario 1 (Figure 3A). The mean coupling percentage was sampled from a normal distribution and the resulting distribution of disruption values followed a very similar bell curve. The same effect occurs with an increase in the number of active reactions – note the mean of the distribution of disrupted yields shifts towards increased disruption with greater reaction activities. In fact, Scenario 2 has between 1.5- and 2-times the number of reactions compared to Scenario 1 (e.g., 34 vs. 17 for pathway 1 and 91 vs. 60 for pathway 2), which leads to a proportional difference in magnitude of disruption observed in the experiments as well, and when the total number of reactions is set to a fixed value as opposed to a percentage, the difference between the two Scenarios almost vanishes (**Figure 4F**).

**Figure 4:**
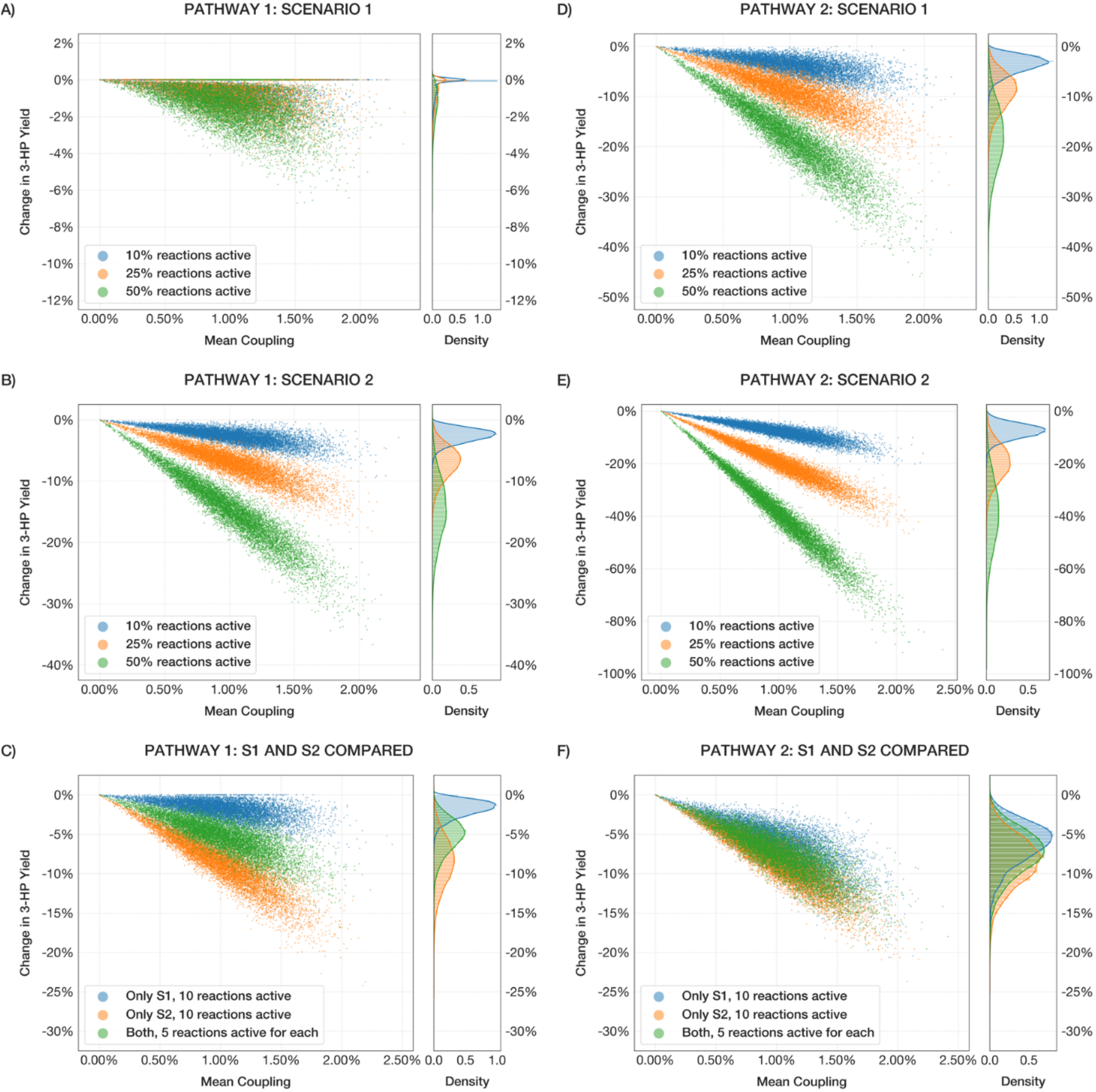
Comparison of simulation runs for the two pathways and both Scenarios, demonstrating the effect of mean coupling percentage and fraction of active promiscuous reactions (10, 25, 50 %) on the yield disruption of 3-HP. (A, D) Pathways 1 and 2 with only Scenario 1-type promiscuous interactions incorporated. (B, E) Pathways 1 and 2 with Scenario 2 interactions. (C, F) Comparison of Scenario 1 (S1) and 2 (S2) interactions for each pathway given a fixed total number of promiscuous reactions. The scatter plots document the results of each individual experiment, while the distribution plots on the right represent the probability density of obtaining a given disruption under the specified conditions.

Scenario 1 for Pathway 1 was an exception to this trend, however. In that instance, only ~17 % of predicted reactions had any effect on 3-HP yield disruption when measured individually – suggesting that only a small fraction of sampled reactions would have a disruptive impact at any given instant under those conditions. As a result, there is a disproportionately large number of instances where promiscuous reactions led to a low magnitude of disruption, particularly for low coupling percentages (**Figure 4A**). The comparison of the two Scenarios for that Pathway further demonstrates the difference in behavior, with Scenario 1 being significantly less disruptive than Scenario 2 on average (**Figure 4C**). This outcome highlights the downside of considering Scenario 1 alone: when few enzymes are being evaluated, the results are highly sensitive to the set of transformation patterns that can be derived from them – subject to knowledge limitations and enzymatic behaviors. Under more realistic biological circumstances, a more diverse range of enzymes participates in metabolism, allowing for more representative results. These observations imply that as promiscuous activity is intensified, it directly competes with 3-HP synthesis pathway flux causing significant disruption. However, such high activity of promiscuous reactions is unlikely under physiological settings, so these exercises likely overestimate the actual extent of disruption.

Overall, Pathway 1 was found to experience comparatively less disruption compared to pathway 2 under the same circumstances, possibly due to a lower number of enzymes, fewer intermediates, and/or lower promiscuous activity of MCR relative to PanD and GabT. In light of our analysis, and considering the similar yield of Pathways 1 and 2, the design of Pathway 1 may be the preferred design option.

## 4. Discussion

In metabolic engineering, a number of strategies are employed to produce a target metabolite of interest, including the introduction of heterologous enzymes and selective overexpression and deletion of certain genes. After such interventions or combinations thereof, it is not uncommon for unexpected and/or undesirable metabolite product profiles to arise from various interactions between the introduced and native machinery in the host. Using *MDFlow*, it is possible to predict such interactions in the form of single- and multi-step pathways enabled by promiscuous enzymatic activity. The method utilizes PROXIMAL to construct reactions arising from promiscuity and relies on FBA to assess their impact on yield or biomass growth rate in pre- and post-modification hosts. The results for the single-gene deletions that were rescued via single- and multi-step enzymatic pathways, which were experimentally validated in prior studies, provide evidence of the ability of *MDFlow* in predicting metabolic network disruption. The results for the 3-HP case illustrate how *MDFlow* can help identify promiscuous interactions early in the design cycle. Further, our sampling-based FBA analysis shows that promiscuity can cause unexpected byproducts and results in yield disruption. Importantly, *MDFlow* can be used to explain byproducts often observed but not well explained in the literature.

Our Scenario 1 and 2 disruption classification has direct correspondence to the network inference classification proposed by Kim et al. [10]. Interference is classified into three groups: those due to (i) heterologous metabolites in new pathways interfering with native metabolism, (ii) native metabolites interfering with a heterologous pathway, and (iii) heterologous pathway intermediates being diverted by promiscuous activity of native enzymes. *MDFlow* identifies the same interactions as long as they’re caused by enzyme substrate promiscuity – with groups (i) and (iii) corresponding to Scenario 2 predictions and group (ii) represented by Scenario 1-type interactions. The computational methodology in *MDFlow* can be further enhanced. It currently uses PROXIMAL to predict promiscuous byproducts; it is possible to use alternatives for promiscuous product prediction. We selected PROXIMAL because we have established confidence in its capabilities in predicting organism-specific enzymatic transformations. Our earlier study of promiscuity using PROXIMAL on non-engineered *E. coli* allowed the discovery of 17 putative enzymatic reactions that explained metabolomics measurement [34]. Regardless of the tool, however, there always remains the issue of false positives. In our work with PROXIMAL, we discarded byproducts that were not documented prior in PubChem, KEGG, or iML1515/1428. Using machine-learning tools that evaluate the likelihood of compound-enzyme interactions, such as SUNDRY [57], might provide further confidence in such predictions. Ultimately, knowing more about the biological sample, such as the concentrations of metabolites and enzymes, and detailed kinetic models [58] can further shed light on the amount of disruption. Importantly, better prediction of enzymatic products and their likelihood can significantly improve the ability to explore complex multi-step scenarios beyond what we presented for our multi-step promiscuity analysis for the Kim et al. study. Another area of improvement is the approach used to determine the direction and maximum flux limits of the predicted reactions. We rely on the preset 1 % average mean coupling percentage to estimate the limits of all reactions, which may not be representative of the higher or lower actual mean coupling percentage under the conditions of a given experiment. Future work may re-evaluate directions and flux limits in the context of each experiment individually. Despite these limitations, the presented results are promising and call for further design exploration of the impact of design promiscuity on engineered microorganisms.

## 5. Conclusion

We presented *MDFlow*, a method for quantitatively evaluating the side effects of engineered modifications on the host metabolic network resulting from enzyme promiscuity. Those side effects can be significant but are difficult to predict in advance, leading to either unexpected behaviors during experiments or failure to take advantage of potentially beneficial interactions. By combining PROXIMAL and FBA in a streamlined workflow, *MDFlow* is capable of both discovering new interactions and evaluating their effects without the need of time-consuming studies of costly *in vivo* experiments. The insights provided by *MDFlow* could be used during the pathway design and optimization process. Multiple pathways for the same target compound could be analyzed and compared using *MDFlow*: when implemented, a pathway with less predicted metabolic disruption may have a higher yield. It is now possible to evaluate disruption effects due to different synthesis pathways and select a synthesis pathway that causes the least disruption.

## Supplementary Files

**Supplementary File 1.** Summary of predicted promiscuous reactions for the multicopy suppressor study done by Patrick et al, categorized by the 8 rescue mechanisms proposed by the authors. For each gene knockout-suppressor pair, we provide the sets of reactions blocked by the deletion and reactions created by the overexpression of the suppressor gene, as well as all relevant growth rates.

**Supplementary File 2.** Predicted two-step serendipitous pathways that may compensate for the deletion of pdxB in *E.coli.* Each pathway is listed in a separate numbered section, with illustrations of all involved metabolites. To view a specific step of a pathway, click on the “Step 1” or “Step 2” link under the section number. To look up a metabolite, template reaction, or an enzyme, click on the corresponding identifier. To view the eQuilibrator query used to compute thermodynamic feasibility for a given pathway, click on the ΔrG’° symbol.

**Supplementary File 3.** Summary of predicted promiscuous reactions for two 3-HP synthesis pathways under Scenario 1 and Scenario 2. For each prediction, we list the enzyme and KEGG template reaction used to derive its biotransformation operator. All metabolites in the model are referred to by their iML1515 identifiers; external metabolites not considered to be a part of normal host metabolism are referenced by their KEGG ids or Pubchem compound identifiers.

## Notes

### Competing Interest Statement

The authors have declared no competing interest.

## References

1. Lee, S.K., et al., Metabolic engineering of microorganisms for biofuels production: from bugs to synthetic biology to fuels. Current opinion in biotechnology, 2008. 2008(6): p. 556–563.

2. Madison, L.L. and G.W. Huisman, Metabolic engineering of poly (3-hydroxyalkanoates): from DNA to plastic. Microbiology and molecular biology reviews, 1999. 1999(1): p. 21–53.

3. Nakamura, C.E. and G.M. Whited, Metabolic engineering for the microbial production of 1, 3-propanediol. Current opinion in biotechnology, 2003. 2003(5): p. 454–459.

4. Trantas, E.A., et al., When plants produce not enough or at all: metabolic engineering of flavonoids in microbial hosts. Frontiers in plant science, 2015. 6: p. 7.

5. George, K.W., et al., Isoprenoid drugs, biofuels, and chemicals—artemisinin, farnesene, and beyond, in Biotechnology of Isoprenoids. 2015, Springer. p. 355–389.

6. Lee, S.Y., et al., Metabolic engineering of microorganisms: general strategies and drug production. Drug Discovery Today, 2009. 14(1-2): p. 78–88.

7. Yoshikuni, Y., et al., Redesigning enzymes based on adaptive evolution for optimal function in synthetic metabolic pathways. Chemistry & Biology, 2008. 2008(6): p. 607–618.

8. Patrick, W.M., et al., Multicopy suppression underpins metabolic evolvability. Mol Biol Evol, 2007. 2007(12): p. 2716–22.

9. Kim, J., et al., Three serendipitous pathways in E. coli can bypass a block in pyridoxal-5’-phosphate synthesis. Mol Syst Biol, 2010. 6: p. 436.

10. Kim, J. and S.D. Copley, Inhibitory cross-talk upon introduction of a new metabolic pathway into an existing metabolic network. Proceedings of the National Academy of Sciences, 2012. 2012(42): p. E2856–E2864.

11. Fujisawa, H., S. Nagata, and H. Misono, Characterization of short-chain dehydrogenase/reductase homologues of Escherichia coli (YdfG) and Saccharomyces cerevisiae (YMR226C). Biochimica et Biophysica Acta (BBA)-Proteins and Proteomics, 2003. 2003(1): p. 89–94.

12. Jessen, H., et al., Compositions and methods for 3-hydroxypropionic acid production. 2015, Google Patents.

13. Inui, M., et al., Expression of Clostridium acetobutylicum butanol synthetic genes in Escherichia coli. Appl Microbiol Biotechnol, 2008. 2008(6): p. 1305–16.

14. Atsumi, S., et al., Metabolic engineering of Escherichia coli for 1-butanol production. Metabolic engineering, 2008. 2008(6): p. 305–311.

15. Nielsen, D.R., et al., Engineering alternative butanol production platforms in heterologous bacteria. Metab Eng, 2009. 11(4-5): p. 262–73.

16. Rathnasingh, C., et al., Production of 3-hydroxypropionic acid via malonyl-CoA pathway using recombinant Escherichia coli strains. Journal of biotechnology, 2012. 2012(4): p. 633–640.

17. Atsumi, S., T. Hanai, and J.C. Liao, Non-fermentative pathways for synthesis of branched-chain higher alcohols as biofuels. nature, 2008. 2008(7174): p. 86.

18. Trinh, C.T., et al., Redesigning Escherichia coli metabolism for anaerobic production of isobutanol. Appl Environ Microbiol, 2011. 2011(14): p. 4894–904.

19. D’ Ari, R. and J. Casadesús, Underground metabolism. BioEssays, 1998. 1998(2): p. 181–186.

20. Nobeli, I., A.D. Favia, and J.M. Thornton, Protein promiscuity and its implications for biotechnology. Nature biotechnology, 2009. 2009(2): p. 157.

21. Khersonsky, O., C. Roodveldt, and D.S. Tawfik, Enzyme promiscuity: evolutionary and mechanistic aspects. Current opinion in chemical biology, 2006. 2006(5): p. 498–508.

22. Tawfik, O.K. and D. S, Enzyme promiscuity: a mechanistic and evolutionary perspective. Annual review of biochemistry, 2010. 79: p. 471–505.

23. Cho, A., et al., Prediction of novel synthetic pathways for the production of desired chemicals. BMC Systems Biology, 2010. 2010(1): p. 35.

24. Campodonico, M.A., et al., Generation of an atlas for commodity chemical production in Escherichia coli and a novel pathway prediction algorithm, GEM-Path. Metab Eng, 2014. 25: p. 140–58.

25. Li, C., et al., Computational discovery of biochemical routes to specialty chemicals. Chemical engineering science, 2004. 59(22-23): p. 5051–5060.

26. Finley, S.D., L.J. Broadbelt, and V. Hatzimanikatis, Computational framework for predictive biodegradation. Biotechnology and bioengineering, 2009. 2009(6): p. 1086–97.

27. Hadadi, N., et al., ATLAS of biochemistry: a repository of all possible biochemical reactions for synthetic biology and metabolic engineering studies. ACS synthetic biology, 2016. 2016(10): p. 1155–1166.

28. Jeffryes, J.G., et al., MINEs: Open access databases of computationally predicted enzyme promiscuity products for untargeted metabolomics. Journal of Cheminformatics, 2015.

29. Li, L., et al., MyCompoundID: using an evidence-based metabolome library for metabolite identification. Anal Chem, 2013. 2013(6): p. 3401–8.

30. Djoumbou-Feunang, Y., et al., BioTransformer: a comprehensive computational tool for small molecule metabolism prediction and metabolite identification. J Cheminform, 2019. 2019(1): p. 2.

31. Amin, S.A., et al., Towards creating an extended metabolic model (EMM) for E. coli using enzyme promiscuity prediction and metabolomics data. Microbial Cell Factories, 2019. 2019(1): p. 109.

32. Hassanpour, N., et al., Biological Filtering and Substrate Promiscuity Prediction for Annotating Untargeted Metabolomics. bioRxiv, 2019: p. 558973.

33. Moriya, Y., et al., PathPred: an enzyme-catalyzed metabolic pathway prediction server. Nucleic acids research, 2010. 38(suppl_2): p. W138–W143.

34. Yousofshahi, M., et al., PROXIMAL: a method for Prediction of Xenobiotic Metabolism. BMC systems biology, 2015. 2015(1): p. 94.

35. Orth, J.D., I. Thiele, and B.Ø. Palsson, What is flux balance analysis? Nature biotechnology, 2010. 2010(3): p. 245.

36. Monk, J.M., et al., iML1515, a knowledgebase that computes Escherichia coli traits. Nat Biotechnol, 2017. 2017(10): p. 904–908.

37. Baba, T., et al., Construction of Escherichia coli K-12 in-frame, single-gene knockout mutants: the Keio collection. Mol Syst Biol, 2006. 2: p. 2006.0008.

38. Chen, Y. and J. Nielsen, Advances in metabolic pathway and strain engineering paving the way for sustainable production of chemical building blocks. Current opinion in biotechnology, 2013. 2013(6): p. 965–972.

39. Della Pina, C., E. Falletta, and M. Rossi, A green approach to chemical building blocks. The case of 3-hydroxypropanoic acid. Green chemistry, 2011. 2011(7): p. 1624–1632.

40. Kumar, V., S. Ashok, and S. Park, Recent advances in biological production of 3-hydroxypropionic acid. Biotechnology advances, 2013. 2013(6): p. 945–961.

41. Schellenberger, J., et al., Quantitative prediction of cellular metabolism with constraint-based models: the COBRA Toolbox v2.0. Nat Protoc, 2011. 2011(9): p. 1290–307.

42. Heller, S.R., et al., InChI, the IUPAC International Chemical Identifier. J Cheminform, 2015. 7: p. 23.

43. RDKit: Open-source cheminformatics. Available from: http://www.rdkit.org.

44. Dahlgren, B., ChemPy: A package useful for chemistry written in Python. The Journal of Open Source Software, 2018.

45. Ye, Q.Z., J. Liu, and C.T. Walsh, p-Aminobenzoate synthesis in Escherichia coli: purification and characterization of PabB as aminodeoxychorismate synthase and enzyme X as aminodeoxychorismate lyase. Proc Natl Acad Sci U S A, 1990. 1990(23): p. 9391–5.

46. Klem, T.J. and V.J. Davisson, Imidazole glycerol phosphate synthase: the glutamine amidotransferase in histidine biosynthesis. Biochemistry, 1993. 1993(19): p. 5177–86.

47. Whalen, W.A. and C.M. Berg, Analysis of an avtA::Mu dl(Ap lac) mutant: metabolic role of transaminase C. J Bacteriol, 1982. 1982(2): p. 739–46.

48. Flamholz, A., et al., eQuilibrator--the biochemical thermodynamics calculator. Nucleic Acids Res, 2012. 40(Database issue): p. D770–5.

49. Noor, E., et al., An integrated open framework for thermodynamics of reactions that combines accuracy and coverage. Bioinformatics, 2012. 2012(15): p. 2037–44.

50. Noor, E., et al., Consistent estimation of Gibbs energy using component contributions. PLoS Comput Biol, 2013. 2013(7): p. e1003098.

51. Noor, E., et al., Pathway thermodynamics highlights kinetic obstacles in central metabolism. PLoS Comput Biol, 2014. 2014(2): p. e1003483.

52. Jiang, X., X. Meng, and M. Xian, Biosynthetic pathways for 3-hydroxypropionic acid production. Applied microbiology and biotechnology, 2009. 2009(6): p. 995–1003.

53. Huang, Y., et al., Co-production of 3-hydroxypropionic acid and 1, 3-propanediol by Klebseilla pneumoniae expressing aldH under microaerobic conditions. Bioresource technology, 2013. 128: p. 505–512.

54. Raj, S.M., et al., Production of 3-hydroxypropionic acid from glycerol by a novel recombinant Escherichia coli BL21 strain. Process Biochemistry, 2008. 2008(12): p. 1440–1446.

55. Kim, K., et al., Enhanced production of 3-hydroxypropionic acid from glycerol by modulation of glycerol metabolism in recombinant Escherichia coli. Bioresource technology, 2014. 156: p. 170–175.

56. Cheng, Z., et al., Enhanced production of 3-hydroxypropionic acid from glucose via malonyl-CoA pathway by engineered Escherichia coli. Bioresource technology, 2016. 200: p. 897–904.

57. Visani, G.M., M.C. Hughes, and S. Hassoun Hierarchical Classification of Enzyme Promiscuity Using Positive, Unlabeled, and Hard Negative Examples. arXiv: 2002.07327 [q-bio.CB], 2020.

58. Khodayari, A. and C.D. Maranas, A genome-scale Escherichia coli kinetic metabolic model k-ecoli457 satisfying flux data for multiple mutant strains. Nature communications, 2016. 7: p. 13806.

