## Supplementary File 2 for "Analysis of Metabolic Network Disruption in Engineered Microbial Hosts due to Enzyme Promiscuity"

Supplementary File 2 — Analysis of Metabolic Network Disruption in Engineered Microbial Hosts due to Enzyme Promiscuity


### Analysis of Metabolic Network Disruption in Engineered Microbial Hosts due to Enzyme Promiscuity

  

Vladimir Porokhin, Department of Computer Science, Tufts University, Medford, MA,

Sara A. Amin, Department of Computer Science, Tufts University, Medford, MA,

Trevor B. Nicks, Department of Chemical and Biological Engineering, Tufts University, Medford, MA,

Venkatesh Endalur Gopinarayanan, Department of Chemical and Biological Engineering, Tufts University, Medford, MA, 

Nikhil U. Nair, Department of Chemical and Biological Engineering, Tufts University, Medford, MA,

Soha Hassoun, Department of Computer Science and Department of Chemical & Biological Engineering, Tufts University, Medford, MA,

  

Supplemenatry File 2

  

|  |  |  |  |
| --- | --- | --- | --- |
| 1 Step 1 Step 2 | C00250    C00002    C00008 | 2.7.6.3  2.7.4.11  →  R03503  R01547   Δ*_r_G'*° = -12.5 kJ/mol | C00018    C00002    C00020 |
| Step 1 | C00250    C00002 | 2.7.6.3  →  R03503 | cid125696    C00020 |
| Step 2 | cid125696    C00008 | 2.7.4.11  →  R01547 | C00018    C00002 |
| 2 Step 1 Step 2 | C00250    C00002    C00008 | 2.7.6.3  2.7.4.12  →  R03503   R00140  R02090  R02094   Δ*_r_G'*° = -12.5 kJ/mol | C00018    C00002    C00020 |
| Step 1 | C00250    C00002 | 2.7.6.3  →  R03503 | cid125696    C00020 |
| Step 2 | cid125696    C00008 | 2.7.4.12  →  R00140  R02090  R02094 | C00018    C00002 |
| 3 Step 1 Step 2 | C00250    C00002    C00008 | 2.7.6.3  2.7.4.14  →  R03503   R00158  R00512  R01665   Δ*_r_G'*° = -12.5 kJ/mol | C00018    C00002    C00020 |
| Step 1 | C00250    C00002 | 2.7.6.3  →  R03503 | cid125696    C00020 |
| Step 2 | cid125696    C00008 | 2.7.4.14  →  R00158  R00512  R01665 | C00018    C00002 |
| 4 Step 1 Step 2 | C00250    C00002    C00008 | 2.7.6.3  2.7.4.16  →  R03503  R00617   Δ*_r_G'*° = -12.5 kJ/mol | C00018    C00002    C00020 |
| Step 1 | C00250    C00002 | 2.7.6.3  →  R03503 | cid125696    C00020 |
| Step 2 | cid125696    C00008 | 2.7.4.16  →  R00617 | C00018    C00002 |
| 5 Step 1 Step 2 | C00250    C00002    C00008 | 2.7.6.3  2.7.4.22  →  R03503  R00158   Δ*_r_G'*° = -12.5 kJ/mol | C00018    C00002    C00020 |
| Step 1 | C00250    C00002 | 2.7.6.3  →  R03503 | cid125696    C00020 |
| Step 2 | cid125696    C00008 | 2.7.4.22  →  R00158 | C00018    C00002 |
| 6 Step 1 Step 2 | C00250    C00002    C00008 | 2.7.6.3  2.7.4.23  →  R03503  R06836   Δ*_r_G'*° = -12.5 kJ/mol | C00018    C00002    C00020 |
| Step 1 | C00250    C00002 | 2.7.6.3  →  R03503 | cid125696    C00020 |
| Step 2 | cid125696    C00008 | 2.7.4.23  →  R06836 | C00018    C00002 |
| 7 Step 1 Step 2 | C00250    C00002    C00008 | 2.7.6.3  2.7.4.25  →  R03503   R00512  R01665   Δ*_r_G'*° = -12.5 kJ/mol | C00018    C00002    C00020 |
| Step 1 | C00250    C00002 | 2.7.6.3  →  R03503 | cid125696    C00020 |
| Step 2 | cid125696    C00008 | 2.7.4.25  →  R00512  R01665 | C00018    C00002 |
| 8 Step 1 Step 2 | C00250    C00002    C00008 | 2.7.6.3  2.7.4.3  →  R03503  R01547   Δ*_r_G'*° = -12.5 kJ/mol | C00018    C00002    C00020 |
| Step 1 | C00250    C00002 | 2.7.6.3  →  R03503 | cid125696    C00020 |
| Step 2 | cid125696    C00008 | 2.7.4.3  →  R01547 | C00018    C00002 |
| 9 Step 1 Step 2 | C00250    C00002    C00008 | 2.7.6.3  2.7.4.4  →  R03503   R00158  R00334  R02098   Δ*_r_G'*° = -12.5 kJ/mol | C00018    C00002    C00020 |
| Step 1 | C00250    C00002 | 2.7.6.3  →  R03503 | cid125696    C00020 |
| Step 2 | cid125696    C00008 | 2.7.4.4  →  R00158  R00334  R02098 | C00018    C00002 |
| 10 Step 1 Step 2 | C06056    C00026    C00081 | 2.6.1.52  2.7.1.1  →  R05085   R02867  R02868   Δ*_r_G'*° = -26.3 kJ/mol | C06054    C00025    C00104 |
| Step 1 | C06056    C00026 | 2.6.1.52  →  R05085 | C21618    C00025 |
| Step 2 | C21618    C00081 | 2.7.1.1  →  R02867  R02868 | C06054    C00104 |
| 11 Step 1 Step 2 | C06056    C00026    C00131 | 2.6.1.52  2.7.1.1  →  R05085   R02867  R02868   Δ*_r_G'*° = -25.8 kJ/mol | C06054    C00025    C00206 |
| Step 1 | C06056    C00026 | 2.6.1.52  →  R05085 | C21618    C00025 |
| Step 2 | C21618    C00131 | 2.7.1.1  →  R02867  R02868 | C06054    C00206 |
| 12 Step 1 Step 2 | C06056    C00002    C00026 | 2.6.1.52  2.7.1.12  →  R05085  R01737   Δ*_r_G'*° = -22.7 kJ/mol | C06054    C00008    C00025 |
| Step 1 | C06056    C00026 | 2.6.1.52  →  R05085 | C21618    C00025 |
| Step 2 | C21618    C00002 | 2.7.1.12  →  R01737 | C06054    C00008 |
| 13 Step 1 Step 2 | C06056    C00002    C00026 | 2.6.1.52  2.7.1.178  →  R05085   R01541  R03387   Δ*_r_G'*° = -22.7 kJ/mol | C06054    C00008    C00025 |
| Step 1 | C06056    C00026 | 2.6.1.52  →  R05085 | C21618    C00025 |
| Step 2 | C21618    C00002 | 2.7.1.178  →  R01541  R03387 | C06054    C00008 |
| 14 Step 1 Step 2 | C06056    C00002    C00026 | 2.6.1.52  2.7.1.30  →  R05085  R00847   Δ*_r_G'*° = -22.7 kJ/mol | C06054    C00008    C00025 |
| Step 1 | C06056    C00026 | 2.6.1.52  →  R05085 | C21618    C00025 |
| Step 2 | C21618    C00002 | 2.7.1.30  →  R00847 | C06054    C00008 |
| 15 Step 1 Step 2 | C06056    C00002    C00026 | 2.6.1.52  2.7.1.45  →  R05085  R01541   Δ*_r_G'*° = -22.7 kJ/mol | C06054    C00008    C00025 |
| Step 1 | C06056    C00026 | 2.6.1.52  →  R05085 | C21618    C00025 |
| Step 2 | C21618    C00002 | 2.7.1.45  →  R01541 | C06054    C00008 |
| 16 Step 1 Step 2 | C06056    C00002    C00026 | 2.6.1.52  2.7.1.53  →  R05085  R07127   Δ*_r_G'*° = -22.7 kJ/mol | C06054    C00008    C00025 |
| Step 1 | C06056    C00026 | 2.6.1.52  →  R05085 | C21618    C00025 |
| Step 2 | C21618    C00002 | 2.7.1.53  →  R07127 | C06054    C00008 |
| 17 Step 1 Step 2 | C06056    C00002    C00026 | 2.6.1.52  2.7.1.58  →  R05085  R03387   Δ*_r_G'*° = -22.7 kJ/mol | C06054    C00008    C00025 |
| Step 1 | C06056    C00026 | 2.6.1.52  →  R05085 | C21618    C00025 |
| Step 2 | C21618    C00002 | 2.7.1.58  →  R03387 | C06054    C00008 |
| 18 Step 1 Step 2 | C00250    C00002    C00008 | 2.7.6.3  2.7.4.7  →  R03503  R04509   Δ*_r_G'*° = -12.5 kJ/mol | C00018    C00002    C00020 |
| Step 1 | C00250    C00002 | 2.7.6.3  →  R03503 | cid125696    C00020 |
| Step 2 | cid125696    C00008 | 2.7.4.7  →  R04509 | C00018    C00002 |
| 19 Step 1 Step 2 | C00250    C00002    C00008 | 2.7.6.3  2.7.4.8  →  R03503   R00332  R02090   Δ*_r_G'*° = -12.5 kJ/mol | C00018    C00002    C00020 |
| Step 1 | C00250    C00002 | 2.7.6.3  →  R03503 | cid125696    C00020 |
| Step 2 | cid125696    C00008 | 2.7.4.8  →  R00332  R02090 | C00018    C00002 |
| 20 Step 1 Step 2 | C00250    C00002    C00008 | 2.7.6.3  2.7.4.9  →  R03503   R02094  R02098   Δ*_r_G'*° = -12.5 kJ/mol | C00018    C00002    C00020 |
| Step 1 | C00250    C00002 | 2.7.6.3  →  R03503 | cid125696    C00020 |
| Step 2 | cid125696    C00008 | 2.7.4.9  →  R02094  R02098 | C00018    C00002 |
| 21 Step 1 Step 2 | C00250    C00001    C00002 | 2.7.6.3  3.6.1.15  →  R03503  R00615   Δ*_r_G'*° = -38.5 kJ/mol | C00018    C00009    C00020 |
| Step 1 | C00250    C00002 | 2.7.6.3  →  R03503 | cid125696    C00020 |
| Step 2 | cid125696    C00001 | 3.6.1.15  →  R00615 | C00018    C00009 |
